# Intravenous colistin use for infections due to multidrug-resistant gram-negative bacilli in critically ill paediatric patients: a systematic review and meta-analysis

**DOI:** 10.1101/465559

**Authors:** Spyridon A. Karageorgos, Hamid Bassiri, George Siakallis, Michael Miligkos, Constantinos Tsioutis

**Affiliations:** Infectious Diseases Working Group, Society of Junior Doctors, Athens, Greece; Division of Infectious Diseases and Center for Childhood Cancer Research, Children’s Hospital of Philadelphia, Perelman School of Medicine at the University of Pennsylvania, USA.; Laboratory of Biomathematics, University of Thessaly School of Medicine, Larissa, Greece; School of Medicine, European University Cyprus, Nicosia, Cyprus

**Keywords:** colistin, multidrug resistance, paediatric, intensive care unit, systematic review.

## Abstract

**Background:** Data are limited regarding the clinical effectiveness and safety of intravenous colistin for treatment of infections by multidrug-resistant gram-negative bacilli (MDR-GNB) in the paediatric intensive care unit (PICU).

**Methods:** Systematic review of intravenous colistin use in critically ill paediatric patients with MDR-GNB infection in PubMed, Scopus and Embase (through January 31^st^, 2018).

**Results:** Out of 1,181 citations, 7 studies were included on the use of intravenous colistin for 405 patients in PICU. Majority of patients were diagnosed with lower respiratory tract infections, with *Acinetobacter baumannii* being the predominant pathogen. Colistin dosages ranged between 2.6-18 mg/kg/day, with none but one case reporting a loading dose. Emergence of colistin-resistance during treatment was reported in two cases. Nephrotoxicity and neurotoxicity were reported in 6.1% and 0.5% respectively, but concomitant medications and severe underlying illness limited our ability to definitively associate use of colistin with nephrotoxicity. Crude mortality was 29.5% (95%CI 21.7-38.1%), whereas infection-related mortality was 16.6% (95%CI 12.2-21.5%).

**Conclusions:** While the reported incidence of adverse events related to colistin were low, reported mortality rates for infections by MDR-GNB in PICU were notable. In addition to severity of disease and comorbidities, inadequate daily dosage and the absence of a loading dose may have contributed to mortality. As the use of colistin for treatment of MDR-GNB infections increases, it is imperative to understand whether optimal dosing of colistin in paediatric patients differs across different age groups. As such, future studies to establish the pharmacokinetic properties of colistin in different paediatric settings are warranted.

## Introduction

The increased incidence of infections due to multidrug-resistant (MDR) or extensively-drug resistant (XDR) gram-negative bacilli (GNB), has a significant impact on patient safety and public health [1,2]. Given the paucity of effective antibiotics that are either available or in clinical development, clinicians have resorted to the use of older agents and combinational therapies for treatment of these infections [3].

Although epidemiological data in paediatric settings are limited, studies worldwide show that the antibiotic resistances of GNB are increasing at an alarming rate. For example, a recent report from the Antibiotic Resistance and Prescribing in European Children (ARPEC) project, demonstrated high rates of resistance to commonly used antibiotic classes in children, especially for *Escherichia coli*, *Klebsiella pneumoniae* and *Pseudomonas aeruginosa* [4], while a national surveillance study among children between 1-17 years of age, revealed a two-fold increase of carbapenem-resistant *P. aeruginosa* in children in the USA from 1999 to 2012 [5]. In another study in European paediatric and neonatal intensive care units (ICU), resistance rates among GNB in intra-abdominal infections were higher than those recorded in non-ICU wards, with MDR rates reaching as high as 13.5% for *P. aeruginosa* and 40.5% for *K. pneumoniae* isolates, whereas the highest MDR rates were noted in Central (24.5% of isolates) and Southeast (11.5% of isolates) Europe [6]. Combinations of antibiotics are often necessary to treat these infections, and in endemic areas these combinations may even be used empirically, prior to confirmation of antibiotic susceptibilities. The limited clinical experience with the use of such treatment strategies in children and the lack of clinical trials involving use of new antibiotics in paediatric populations [7] further aggravate the situation.

Colistin, which was discovered in 1949 and used since the 1950s has found a recent resurgence in use for treatment of infections by MDR- and XDR-GNB [8]. Although the dosing and safety profile of colistin have been extensively studied in adults, several issues remain unresolved in paediatric patients, particularly the pharmacokinetics and optimal dosing of children of different ages [8].

The aim of this systematic review was to evaluate the available evidence concerning the clinical effectiveness and safety of colistin in the treatment of infections due to MDR-GNB in the PICU.

## Material and methods

### Data Sources and Search

This review adopted the Preferred Reporting Items for Systematic Reviews and Meta-Analyses (PRISMA) guidelines [9]. We performed a literature search in PubMed, Scopus and Embase databases from 2000 through 2018 (last day of search was January 31^st^, 2018). The search term algorithm applied in PubMed was: (colistin OR polymyxin E) AND (child* OR pediatric* OR paediatric* OR toddler* OR adolescent*); whereas in Scopus and EMBASE the algorithm was: (colistin OR polymyxin B) AND (child OR children). The references of articles found in this manner were subsequently also searched for relevant articles that could meet the PRISMA criteria.

### Study selection

Articles were eligible for inclusion, provided they fulfilled the following criteria: inclusion of at least 5 paediatric (defined as ≥1 month and ≤18 years of age) patients, and the use of intravenous colistin for treatment of infections due to MDR, XDR or pandrug-resistant GNB in a PICU setting. According to published definitions, a multidrug-resistant (MDR) pathogen is one that is resistant to at least 3 antibiotic classes, an extensively-drug-resistant (XDR) pathogen is resistant to all except one or two antibiotic classes, and a pandrug-resistant (PDR) pathogen is resistant to all antibiotic classes [1]. The following studies were excluded from analysis: studies conducted exclusively in patients with cystic fibrosis; studies that did not include colistin treatment outcomes; studies that included only neonates or were performed in a neonatal ICU; studies performed outside of the PICU; studies that used colistin for treatment of diarrhoea, for decontamination, or for prophylaxis; studies that used polymyxin B or polymyxin E formulations; studies exclusively concerning topical, oral, intraventricular, or intrathecal administration of colistin; conference abstracts, letters to the editor, articles without new data and reviews, and studies published in languages other than English. Authors of included studies were contacted for further clarifications when needed.

### Outcomes and definitions

The primary outcome measure was all-cause mortality in patients who received intravenous colistin for treatment of the MDR- or XDR- or PDR-GNB infection. If available, infection-related mortality was also recorded. All reported outcome measures were classified according to the definitions provided by each study. The secondary outcomes of interest included the following: clinical cure/improvement; microbiological eradication of the original pathogen in a subsequent culture; number and type of adverse events which were either directly related or possibly related to colistin use, per the determination of the authors; and development of resistance to colistin during treatment. Nephrotoxicity and neurotoxicity were based on definitions used by each included study. The quality of the evidence regarding outcomes was assessed using the Grading of Recommendations Assessment, Development and Evaluation (GRADE) algorithm [10].

### Data extraction

Two investigators (CT and SAK) independently reviewed the titles and abstracts of the citations for potentially relevant articles using Abstrackr [11]; the full text publications of potentially relevant articles were retrieved and rescreened by the same two investigators. Disagreements were resolved by consensus with a third author (MM). Data were extracted by SAK and GS, using Excel^®^ and included author, year, type of study, geographic region where study was conducted, number and characteristics of patients, their underlying diseases, severity of disease, type of infection, causative agents, site of pathogen isolation, colistin dosage, use of other forms of colistin (intraventricular/intrathecal, inhalation/nebulised, oral, topical), concomitant antibiotics, and outcomes as defined above.

### Statistical analysis

We calculated the summary mortality rate and corresponding 95% confidence interval (CI), using the random-effects model with arcsine transformation for proportions [12]. We assessed statistical heterogeneity using the I2 statistic [13]. Statistical analysis was performed with OpenMetaAnalyst (http://www.cebm.brown.edu/open_meta/).

## Results

### Literature search

For this systematic review (Fig.1), we screened 1,181 non-duplicate citations; we excluded 1,030 as irrelevant and 151 articles were retrieved for full-text review. Study selection process is presented graphically in Figure 1. Of these, 7 studies met our inclusion criteria [14–20]. In one study, authors kindly provided clarifications and additional data [15]. All studies were retrospective and published between 2009 and 2018. Five studies were conducted in Asia [16–20] and two were conducted in Europe [14,15]. The majority of included studies were single arm retrospective cohorts; as a result the overall quality of the evidence that contributed to our systematic review was rated as low to very low [10].

**Figure 1.**
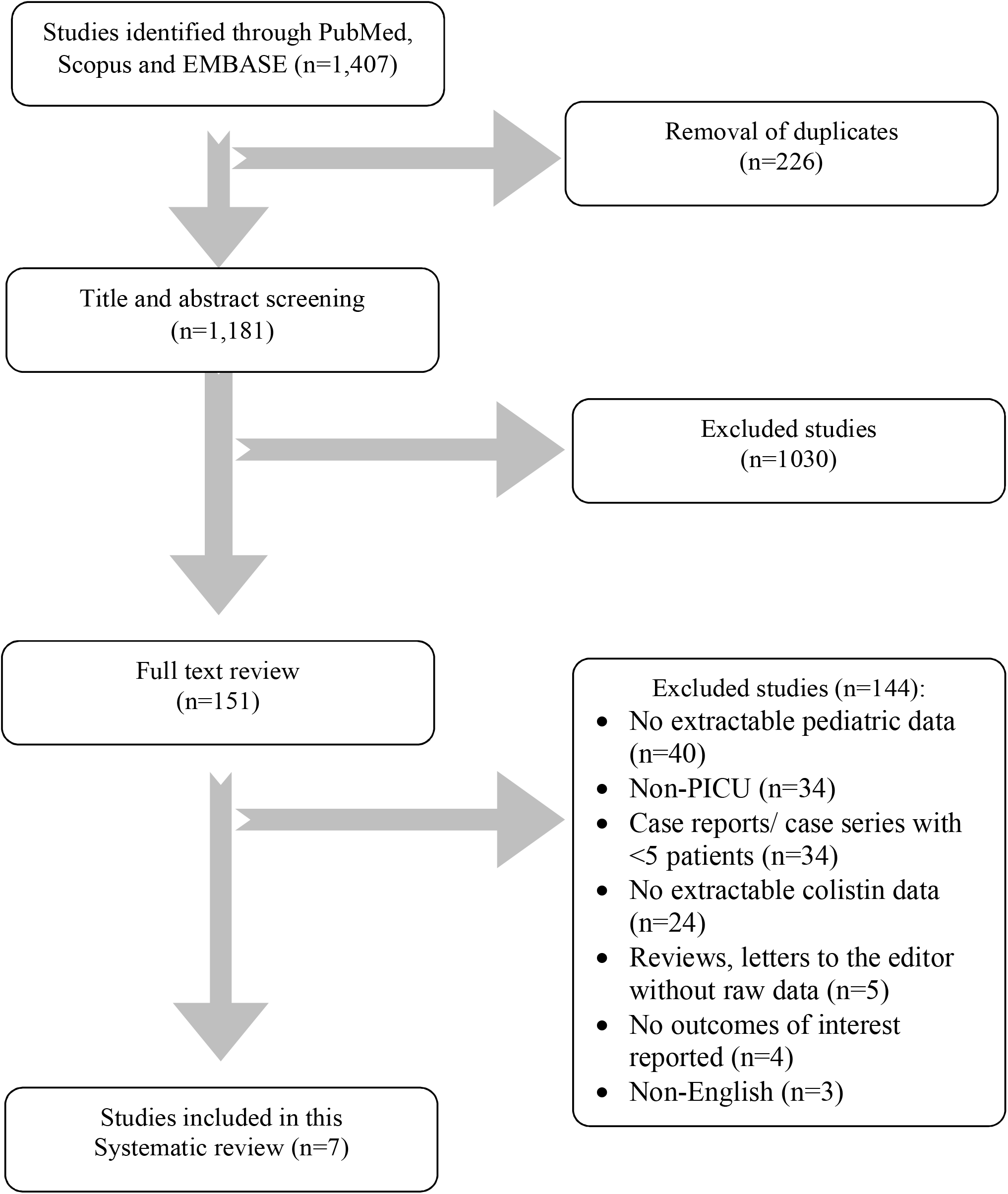
Study selection.

### Study characteristics

Data were available for 405 patients who received IV colistin for treatment of infections due to MDR-GNB [14–20]. Sixty one percent (248/405) of patients were male. The age of patients ranged from 1 month to 18 years. Data on patient demographics, underlying disorders, type of infections that warranted the initiation of colistin therapy, isolated microorganisms, site of pathogen isolation, and treatment, are presented in Table 1.

**Table 1.** Study Characteristics.

| **Study, Country, Year [Ref]** | **No. patients (pts), or courses (crs) [male]** | **Mean age<sup>a</sup> (range)** | **Underlying disorders (no. patients)** | **Type of infection as reported by authors (no. patients)** | **Causative organism (no. patients)<sup>b,c</sup>** | **Site of pathogen isolation** | **Type of colistin formulation (add'l mode of delivery)** | **Dose of colistin and duration of treatment<sup>a</sup>** | **Concomitant therapeutic regimens (n patients)** |
| --- | --- | --- | --- | --- | --- | --- | --- | --- | --- |
| Falagas et al Greece 2009 | 6 pts [5] | 11y (14m-13y) | GM1 gangliosidosis (1)<br>Cerebral palsy (1)<br>None (4) | LRTI (4)<br>BSI (1)<br>LRTI+BSI (1) | PA (3)<br>AB (2)<br>KP+AB (1) | Bronchial secretions (4)<br>Blood (1)<br>Blood & bronchial secretions (1) | Colistimethate sodium (ND) | 5mg/kg/d ÷3 doses<br>Mean 10.7d | Piperacillin/tazobactam (1)<br>Gentamicin (1)<br>Imipenem/silastatin (1)<br>Liposomal amphotericin B (1)<br>Metronidazole (1)<br>Vancomycin (1) |
| Iosifidis et al Greece 2010 | 12 pts 18 crs [5] | 5y (1.5m-14y) | RDS, psychomotor retardation (2)<br>Trauma (2)<br>Hydrocephalus, EVD (1)<br>TB meningitis, EVD (1)<br>Meningomyelocele, hydrocephalus, EVD | RTI(4)<br>CNS infection (2)<br>BSI+RTI (2)<br>BSI (1)<br>Trauma (1)<br>Trauma+ RTI (1) | AB (6)<br>PA (1)<br>ECI (1)<br>AB+SM (1)<br>AB+KP (1)<br>KP+PA+SM (1) | Bronchial secretions (4)<br>Blood & bronchial secretions (2)<br>CSF (2)<br>CSF and | Colistimethate sodium (3 IVT; 2 inhaled) | Range 40,000-225,000 IU/Kg/d<br>Range 7-133d (3/12 rec'd < | Aminoglycosides (16/18 crs)<br>Carbapenems (12/18 crs)<br>Teicoplanin (5/18 crs)<br>Vancomycin (4/18 crs)<br>Metronidazole (4/18 crs)<br>Fluconazole (4/18 crs)<br>Amphotericin B (4/18 crs) |
|  |  |  | (1)<br>Spinal tumor (1)<br>Brain tumor+<br>leukemia (1)<br>RDS+ seizures (1)<br>Wilson disease+ liver<br>transplantation (1)<br>Respiratory failure+<br>obesity (1) | CNS<br>infection+<br>RTI (1) | AB+KP+<br>PA<br>+ECI(1) | bronchial<br>secretions<br>(2)<br>Trauma and<br>bronchial<br>secretions<br>(1)<br>Trauma (1) |  | 21d) | TMP/SMX (3/18 crs)<br>Piperacillin/tazobactam<br>(2/18 crs)<br>Voriconazole (1/18 crs)<br>Tigecycline (1/18 crs)<br>Cloxacillin (1/18 crs)<br>Clarithromycin (1/18<br>crs)<br>Gancyclovir (1/18 crs)<br>Anti-TB (1/18 crs) |
| Kapoor et al<br>India<br>2013 | 50 pts<br>[30] | 36m<br>(1m-12y) | Septic shock (29)<br>Ileal perforation (3)<br>Esophageal stricture<br>(1) | CAP (18)<br>VAP (12) | AB (35)<br>PA (9)<br>KP (7)<br>ECo (3)<br>ECI (1) | Endotrachea<br>l lavage (25)<br>Blood (22)<br>Urine (3)<br>Pleural fluid<br>(1)<br>5 pts with<br>pos. cultures<br>from >1 site<br>with same<br>organism | Colistin<br>NOS | 50,000-75,000<br>IU/Kg/d÷<br>3 doses<br>Mean<br>14.3d<br>(range 7-21). | Meropenem (35)<br>Piperacillin/sulbactam<br>(10)<br>Vancomycin (6)<br>Fluconazole (4)<br>Amphotericin B (2) |
| Paksu et al<br>Turkey<br>2012 | 79 pts<br>87 crs<br>[43] | 30m | Chronic<br>neurological/neuromu<br>sclar disease (26)<br>Congenital heart<br>disease (10)<br>Primary<br>immunodeficiency (8)<br>Metabolic disorders | VAP (49)<br>VAP+BSI<br>(18)<br>BSI (9)<br>Sepsis (8)<br>CNS<br>infection (1)<br>Soft tissue | AB<br>(52/87)<br>PA<br>(16/87)<br>KP (1/87)<br>AB+PA+<br>KP (7/87)<br>NR | Tracheal<br>aspirate<br>fluid (63/87)<br>Blood<br>(28/87)<br>Skin swabs,<br>conjunctival<br>swabs (5/87) | Colistin<br>NOS (1<br>IVT) | Mean±SD<br>5.4±0.6<br>mg/Kg/d<br>Mean<br>17.2±8.4d<br>(2-62) | Glycopeptides (36)<br>Antifungal agents (32)<br>Carbapenems (28)<br>Aminoglycosides (27)<br>Fluoroquinolones (12)<br>Linezolid (12)<br>Cefoperazone/sulbactam<br>(8) |
|  |  |  | (7)<br>Malignancy (2)<br>Others (9) | infection (1)<br>Peritonitis (1) | (11/87) | Others (peritoneal, CSF) (2/87) |  |  | Piperacillin/tazobactam (6)<br>Antiviral agents (5)<br>Others (8) |
| Phan et al<br>Vietnam<br>2014 | 104<br>[73] | 4m | NR | VAP (45) | AB, KP, PA consisted 92% of organisms | NR | Colistimethate sodium (NR) | Mean 6.2±1.2 mg/Kg/d<br>Median 17d (IQR 11-23) | NR |
| Polat et al<br>Turkey<br>2015<sup>d</sup> | 32<br>[15]<br>18<br>[7] | 18m<br>13.5m | Metabolic disorders (16)<br>Chronic neurological/neuromuscular diseases (15)<br>Malignancy (9)<br>Chronic liver disease, Congenital heart disease and thromboembolic events (6)<br>Primary immunodeficiency (4) | VAP 50/50 | AB (37)<br>PA (13) |  | Colistimethate sodium (0)<br>Colistimethate sodium (18 aerosolized) | 3.2mg/kg/d (2.6-5 mg/kg/d). 16 days (10-22)<br>3.4mg/kg/d (2.8-5 mg/kg/d). 14 days (5-21) | Glycopeptides (33/50)<br>Carbapenems (30/50)<br>Aminoglycosides (21/50)<br>Cefoperazone/sulbactam (9/50)<br>Piperacillin/tazobactam (8/50)<br>Fluoroquinolones (5/50) |
| Sahbudak Bal et al<br>Turkey<br>2017 | 104<br>[70] | 55.9m | Chronic neurological/neuromuscular disorder (27)<br>Congenital heart disease (14) | VAP (60)<br>CLABSI+ sepsis (22)<br>CLABSI+ VAP (7/) | AB (57/104)<br>PA (25/104)<br>KP | Endotracheal lavage 59<br>Blood 26<br>Urine 10<br>CSF 3 | Colistin NOS (13 intrathecal) | 5 mg/kg/day in 3 doses<br>12.5±6.4 | Aminoglycosides 76/104<br>Glycopeptides 33/104<br>Amphotericin B 30/104<br>Meropenem 7/104<br>Antiviral agents |
|  |  |  | Chronic lung disease (11)<br>Cancer (10)<br>BM/solid transplantation (10)<br>Primary immune deficiency (6)<br>Chronic liver disease (6)<br>Others (3) | CLABSI+<br>UTI (4)<br>VAP+ UTI (3)<br>UTI (3)<br>Shunt infection (3)<br>SSI (2) | (12/104)<br>AB+KP+<br>PA (3/104)<br>)<br>ECo<br>(2/104)<br>ECI<br>(1/104) | Peritoneal fluid 2 |  | (2-30)<br>(mean±SD (range) | (aciclovir, ganciclovir) 6/104<br>Ciprofloxacin 5<br>TMP-SMX 2/104 |

AB=*A. baumannii*; BSI= Blood stream infection; CAP= community acquired pneumonia; CLABSI=Central line-associated bloodstream infection; CNS= Central nervous system; CSF= Cerebrospinal fluid; ECI= *E. coli*; ECo=*E. cloacae*; EVD= External ventricular device; IU= International units; IVT=Intraventricular; KP= *K. pneumoniae*; LRTI=Lower respiratory tract infection; m=months; ND=No data; NOS=not otherwise specified; NR= Not reported; PA=*P. aeruginosa*; PICU=Pediatric Intensive Care Unit; RDS= acute respiratory distress syndrome; RTI= Respiratory tract infection; SD=standard deviation; SM=*S. maltophilia*; SSI= Surgical site infection; TB=tuberculosis; TMP-SMX- trimethoprim-sulfamethoxazole; UTI=Urinary tract infection; VAP=Ventilator-associated pneumonia; y=years.
<sup>a</sup> Data are provided as medians (range) unless otherwise specified.
<sup>b</sup> Some infections were polymicrobial.
<sup>c</sup> All presented data refer to MDR GNB unless otherwise specified.
<sup>d</sup> Two patients were not admitted to the PICU during colistin administration

### Type of infections and isolated pathogens

The most common infections that warranted use of colistin were lower respiratory tract infections (primarily ventilator-associated pneumonia), followed by bloodstream infection, urinary tract infection, central nervous system infection (including external ventricular drainage-associated ventriculitis or meningitis), and wound infection; sites from which these GNB were isolated are presented in Table 1. The most commonly isolated pathogen was *Acinetobacter baumannii*, followed by *P. aeruginosa* and *K. pneumoniae*. Other reported microorganisms were *Enterobacter cloacae, E. coli* and *Stenotrophomonas maltophilia*. The exact number of polymicrobial infections could not be estimated, as this was not clarified in several of the studies.

### Colistin treatment

Concerning colistin dosage, 4 studies reported colistin dose in milligrams, ranging from 2.6 to >9 mg/kg/day [14,17–20], two studies [15,16] reported colistin dose in international units (IU), ranging from 40,000 to 225,000 IU/kg/day (estimated at 3.2-18 mg/kg/day according to Ortwine et al [21]). Three studies reported that colistin was administered in 3 divided daily doses [14,16,20]. A loading dose was reported in one patient [15], consisting of 225,000 IU/kg (estimated at 18 mg/kg [21]). Mean duration of colistin treatment was 10.8 to 31.6 days (range 2-133 days). In the majority of included studies, colistin was used in combination therapy, most commonly with carbapenems, glycopeptides, and aminoglycosides [14–17,19,20]. Three studies reported co-administration of IV colistin with other formulations of colistin (aerosolised, intraventricular) as presented in Table 1 [15,17,19].

### Mortality

Mortality was reported in all included studies (405 patients) [14–20]. The summary all-cause mortality was 29.5% (95% CI 21.7-38.1%, I^2^=64.7%; Figure 2). In 6 studies that reported an association between mortality and infection [14–17,19,20], the infection-related mortality was 16.6% (95% CI 12.2-21.5%, I^2^=13.1%,; Figure 3).

**Figure 2.**
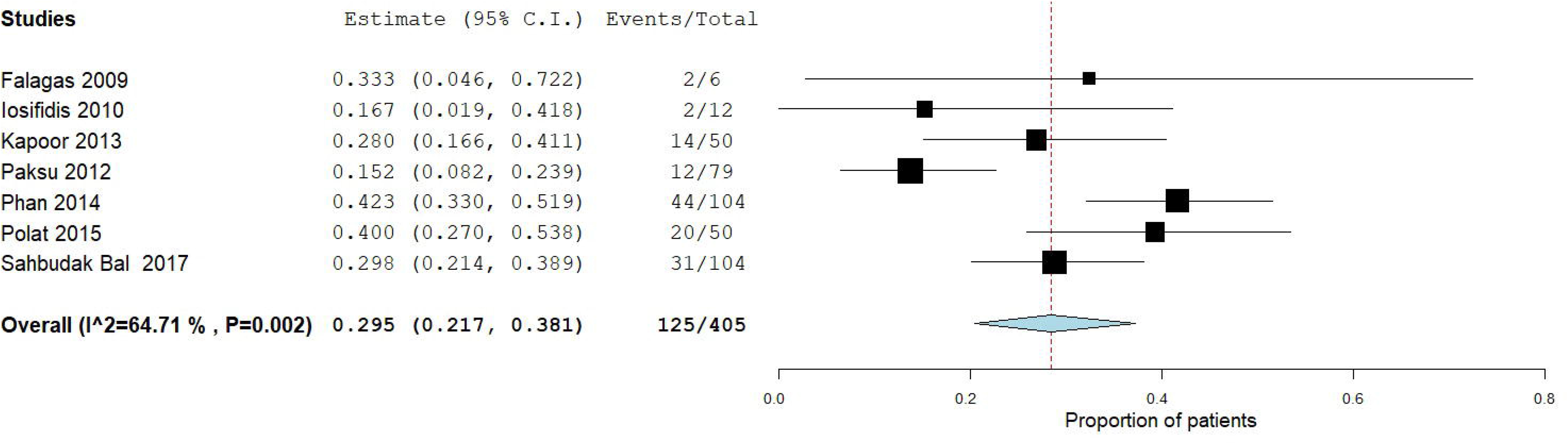
All-cause mortality.

**Figure 3.**
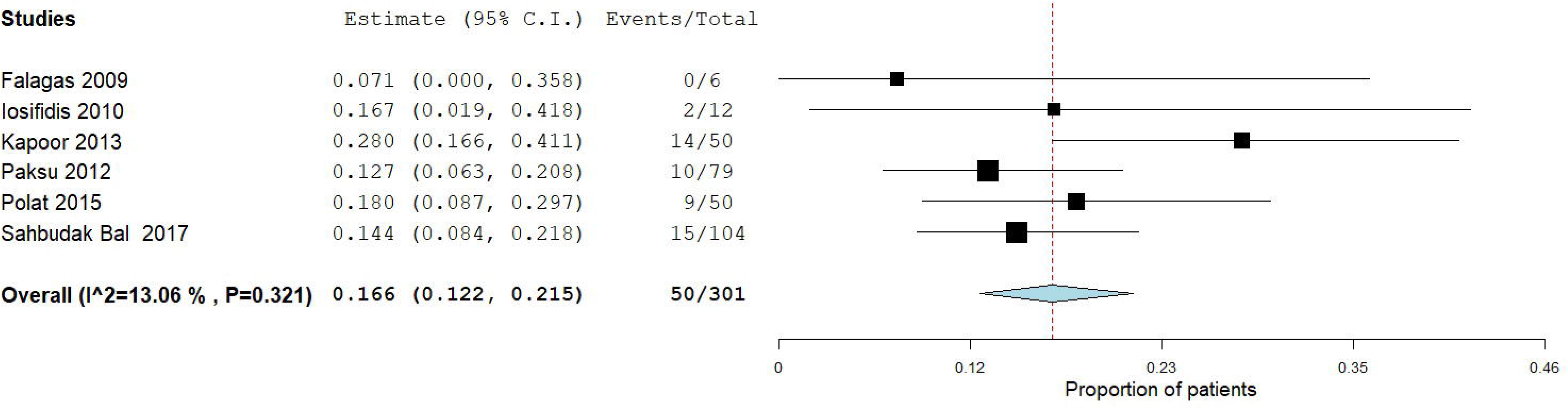
Infection-related mortality.

### Clinical outcomes, microbiological eradication, and adverse events

All studies provided information on secondary outcomes (Table 2). The summary clinical cure/improvement, as defined by the study authors in 6 studies [14–16,18–20] was 73.1% (95% CI 64.4-81.0%, I^2^=58.2; Figure 4). In one study, clinical cure/improvement was reported in 70/87 (80.4%) episodes [17].

**Table 2.** Outcomes.

| **Study, Year [Ref]** | **No. patients** | **Clinical Cure/improvement** | **Definition of clinical cure/improvement** | **Microbiological eradication** | **Definition of microbiological eradication** | **Deaths (related to infection that required colistin therapy)** |
| --- | --- | --- | --- | --- | --- | --- |
| Falagas et al 2009 | 6 | 6 | Improvement of symptoms and signs of the index infection and the laboratory values | 6 | Eradication of the pathogen isolated initially by culture | 2 |
| Iosifidis et al 2010 | 12 (18 courses) | 16/18 courses<br>10/12 patients | No clinical or laboratory signs of infection | ND | ND | 2 (2 related to index infection) |
| Kapoor et al 2013 | 50 | 36 | Resolving presenting signs and symptoms | 44/46 with available follow up cultures | Absence of the same organism from the same site on follow up cultures | 14 (all had MODS) |
| Paksu et al 2012<sup>a</sup> | 87 episodes (79 patients) | 70 episodes | Complete recovery from clinical findings of index infection at end of colistin treatment | 68 episodes | Eradication of the causative microorganism in the final culture | 12 (10 related to index infection) |
| Phan et al 2014 | 104 | 61 | ND | ND | ND | 44 (ND) |
| Polat et al 2015<sup>b</sup> | 50 | 38 | Clinical cure or improvement | 38 | Bacterial eradication (no growth of the causative organism on follow-up cultures regardless of clinical outcome) | 20 (9 related to index infection) |
| Sahbudak Bal et al 2017 | 104 | 79 | Complete recovery from clinical findings of index<br>infection | 62 | Culture clearance at the end of therapy | 31 (15 related to index infection) |

<sup>a</sup> Numbers provided in episodes.
MODS= Multiple organ dysfunction syndrome; ND= No data

**Figure 4.**
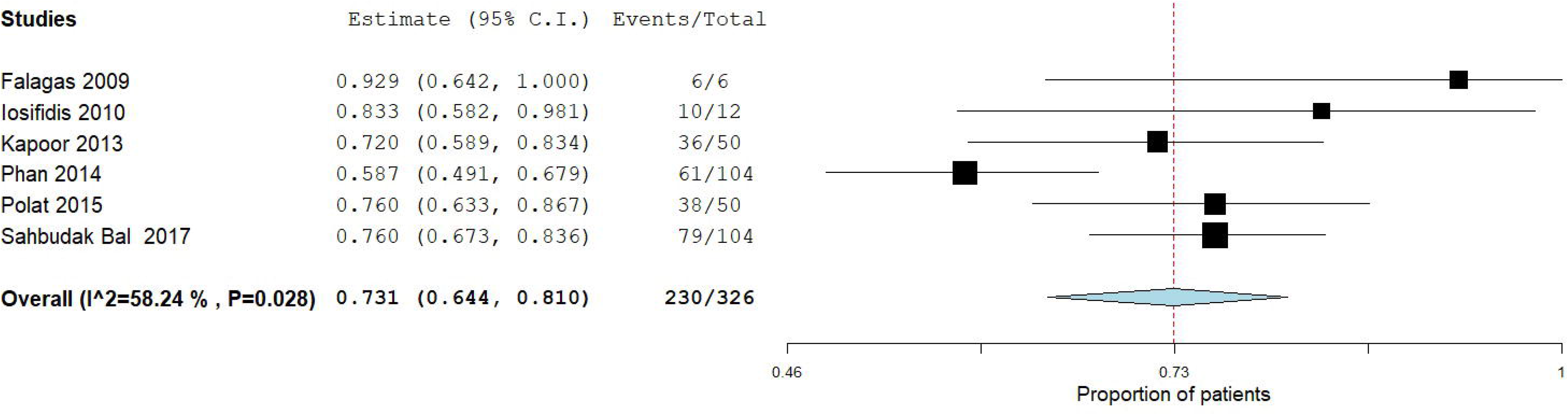
Clinical cure/improvement.

In 5 studies that included data on follow-up cultures, microbiological eradication was confirmed in 150/206 (72.8%) of patients [14,16,19,20] and in 68/87 (78.2%) of episodes [17]. Development of resistance to colistin during therapy was reported in 2 patients, after prolonged use of intravenous and intraventricular colistin in one patient, and intravenous and aerosolised colistin in the other [15].

Data regarding adverse events were reported in all seven studies (Table 3) [14–20]. Nephrotoxicity (defined either as creatinine elevation from baseline or decrease of creatinine clearance during colistin treatment) was detected in 25/405 (6.1%) of patients. Notably, all patients with renal injury were either receiving concomitant nephrotoxic agents (such as vancomycin, aminoglycosides, radio contrast, or amphotericin B), or had pre-renal impairment or multiorgan failure. Management of renal impairment was reported in 5 patients as follows: colistin dose was adjusted in three [15,16], colistin was discontinued in one [17], and colistin was continued with renal replacement therapy in another [17]. In four patients [15,16,19] renal function returned to normal after colistin was adjusted or stopped, while three patients died of multiorgan failure [16]; the outcome of renal function was not reported in the remaining patients. Neurotoxicity was detected in 2/405 (0.5%) of patients [17]. Neurotoxicity was described as generalised tonic-clonic seizures on the first day of colistin administration without recurrence thereafter. Finally, in one study [16], four cases of microscopic haematuria in the context of disseminated intravascular coagulation were also reported.

**Table 3.** Adverse events.

| **Study, Year [Ref]** | **No. patients** | **Patients with nephrotoxicity** | **Definition of nephrotoxicity** | **Patients with neurotoxicity** | **Definition of neurotoxicity** | **Patients with other adverse events** |
| --- | --- | --- | --- | --- | --- | --- |
| Falagas et al 2009 | 6 | 0 | 2-fold increase in creatinine from baseline to >1.3 mg/dL | 0 | Level of consciousness, seizures, visual disturbance, neuromuscular blockade | 0 |
| Iosifidis et al 2010 | 12 (18 courses) | 1 | Elevation of creatinine values beyond the estimated normal range for age | 0 | Neuromuscular blockade, seizures, disturbance of consciousness | 0 |
| Kapoor et al 2013 | 50 | 5* (3 with MODS, 2 with concomitant vancomycin) (nephrotoxicity at 3<sup>rd</sup>-6<sup>th</sup> day of treatment) | 2-fold increase in creatinine from baseline or a 30% decrease in creatinine clearance | 0 | Level of consciousness, seizures, visual disturbance, neuromuscular blockade | 4 (microscopic hematuria in the context of DIC) |
| Paksu et al 2012 | 79 (87 episodes) | 2 (both with concomitant gentamicin) (1 developed at day 8) | Serum creatinine >1.1 mg/dL or a 50% reduction in creatinine clearance or need for renal replacement therapy at any time | 2 (tonic-clonic seizures) | Neuromuscular blockade, seizures, change in level of consciousness | 0 |
| Phan et al 2014 | 104 | 5 (all with MODS) | ND | 0 | ND | 0 |
| Polat et al 2015 | 50 | 1* (concomitant vancomycin & radiocontrast) (day 8) | Increase in serum creatinine by $\geq 50$ % from the baseline and/or elevation of serum creatinine beyond the estimated normal range for age | 0 | ND | ND |
| Sahbudak Bal et al | 104 | 11 (all with concomitant | Blood creatinine level >1.2 | 0 | Paresthesia, neuromuscular | 0 |
| 2017 |  | nephrotoxic agents)<br>(4 at day 0-3<br>6 at day 3-7<br>1 at day 7-14) | mg/dL or an increase of 50% above baseline creatinine or a decline in renal function |  | blockade, seizures, change in level of consciousness |  |

\*: Not clearly related to colistin use.
DIC= Disseminated intravascular coagulation; MODS= Multiple organ dysfunction syndrome;
ND= No data.

## Discussion

MDR-GNB have emerged as a significant cause of infection in paediatric settings, resulting in increased morbidity and mortality [2,3]. Colistin, to which many of these MDR-GNB retain sensitivity, has gained increased use over the last decade [8], a fact that is supported by the observation that all included studies were published after 2009. In this systematic review, we found that colistin use had a relatively low rate of adverse events and resulted in favourable clinical outcome in 73.1% of paediatric patients with MDR-GNB infections hospitalised in a PICU setting, although the pooled and infection-related mortality rates were 29.5% and 16.6% respectively. It is also worth noting that the majority of included studies were conducted in countries that are known to have a high prevalence of MDR-GNB, signifying the importance of prudent use of colistin, to preserve its efficacy against these pathogens.

Our results demonstrate that the majority of MDR-GNB infections in the PICU treated with colistin were lower respiratory tract infections (and especially ventilator-associated pneumonia). Furthermore, we show that *A. baumannii* was the predominant pathogen, and most treated patients possessed significant comorbidities. Nonetheless, colistin achieved clinical improvement/cure and microbiological eradication in ~70% of infections. The observed pooled mortality rate in our study reflects that of critically ill children with severe infections observed in other multicentre [22,23] and single-centre [24–26] studies. Yet a previous systematic review of colistin use in children reported a lower crude mortality rate (7.4%) [27]. There are however, critical differences between these two studies: in the other study, not all patients were critically ill, MDR pathogens were infrequent, a proportion of patients were receiving systemic colistin for prophylaxis rather than treatment, and the majority of included studies were case reports. In support of our contention that these features may dramatically alter the mortality rates, adults treated with colistin for infections by carbapenem-resistant GNB had rates of clinical cure/improvement and mortality that were more comparable to what we report, with estimated pooled mortality rates between 33.8% - 35.7% [28,29].

As far as adverse events are concerned, colistin demonstrated a relatively good rate of tolerability. The most frequently reported adverse event was nephrotoxicity, detected in a small proportion of paediatric patients (6.1%). However, a clear causal association with colistin could not be ascertained, since most of these patients received concomitant nephrotoxic agents or had underlying conditions that compromised renal function. These are factors that have been previously associated with colistin-related nephrotoxicity in adults [30]. Of note, the reversibility of kidney injury after cessation or dose adjustment of colistin, supports the necessity for continuous awareness and frequent monitoring of renal function. Neurotoxicity complicated only 0.5% of paediatric patients, and other adverse events were reported in a small proportion. We believe that the lower rates of acute kidney injury and neurotoxicity [27] reported in the previous systematic review of colistin use in children are again attributable to the differences in the study characteristics. However, it is worthy to note that the adverse event profile of colistin use in adults is different; in a recent meta-analysis regarding the use of colistin in MDR-GNB infections, colistin-related nephrotoxicity was noted to be much higher (19.2%), while neurotoxicity was not reported [28].

The notable difference in rates of nephrotoxicity highlights the observation that during childhood, various physiologic alterations affect drug pharmacokinetics (e.g. volume of distribution, protein binding and renal clearance) and thus, conclusions derived from adult studies do not readily apply to paediatric populations [31], thus highlighting the need for future studies regarding the pharmacodynamics of colistin therapy in paediatric patients. Indeed, the need for a loading dose and the optimal dosing of colistin is likely to vary between different age groups according to renal function and body weight, or body surface area [31]. In support of this notion, adult cases of nephrotoxicity have been associated with excessive colistin dosage, owed to calculation of doses based on ideal body weight rather than actual body weight [32]. Finally, future paediatric studies should employ strict definitions regarding colistin-related clinical outcomes and adverse events, in order to better evaluate the side effect profile of colistin and these studies should report all potential colistin-related side effects instead of focusing primarily on the historically reported ones (nephrotoxicity and neurotoxicity).

In closing, we would like to acknowledge two limitations of our study. Firstly, the studies reported herein were all cohort studies; while this diminishes the quality of evidence, we also cannot exclude an outcome reporting bias favouring the reporting of successful treatments. Secondly, in one study [17] infection episodes were reported rather than infected patients, making interpretations from this study difficult to correlate to the number of affected children. Despite its shortcomings, our study argues that colistin may be relatively safe to use in children with severe MDR-GNB infections in which therapeutic options are limited.

## Conclusions

In an era of increasing antimicrobial resistance, the present systematic review suggests that colistin for infections by MDR-GNB in paediatric patients in the PICU results in favourable clinical outcomes and has an acceptable safety profile. Until a more robust understanding of the pharmacokinetics and pharmacodynamics of colistin in various paediatric populations has been completed, colistin dosing should be carefully estimated in order to achieve maximum efficacy with the lowest possibility of adverse events, as this drug remains one of the last resorts for treatment of infections by MDR-GNB.

## Ethical approval

Not required.

## Transparency declarations

none to declare.

## Author Contributions

Spyridon A. Karageorgos (SAK): conceptualised and designed the study, participated in data acquisition, extraction and interpretation, prepared tables, wrote and drafted the initial manuscript and approved the final manuscript as submitted.

Hamid Bassiri (HB): participated in data interpretation, reviewed and revised the manuscript and approved the final manuscript as submitted.

George Siakallis (GS): participated in data extraction and interpretation, reviewed and revised the initial manuscript and approved the final manuscript as submitted.

Michael Miligkos (MM): participated in data analysis and interpretation, reviewed and revised the initial manuscript and approved the final manuscript as submitted.

Constantinos Tsioutis (CT): conceptualised and designed the study, interpreted the data, wrote and drafted the initial manuscript, reviewed and revised the manuscript and approved the final manuscript as submitted.

